# Imaging Electrical Activity of Retinal Ganglion Cells with Fluorescent Voltage and Calcium Indicator Proteins in Retinal Degenerative *rd1* Blind Mice

**DOI:** 10.1101/2023.12.10.571014

**Authors:** Younginha Jung, Sungmoo Lee, Jun Kyu Rhee, Chae-Eun Lee, Bradley J. Baker, Yoon-Kyu Song

**Affiliations:** Graduate School of Convergence Science and Technology, Seoul National University, Seoul, 08826, South Korea; Brain Science Institute, Korea Institute of Science and Technology, Seoul, 02792, South Korea; Division of Bio-Medical Science and Technology, Korea University of Science and Technology, Seoul, 02792, South Korea

**Keywords:** Retinal degeneration, Retinitis pigmentosa, Retinal ganglion cells, Genetically-encoded voltage indicator, Genetically-encoded calcium indicator, Epi-retinal stimulation

## Abstract

In order to understand the retinal network, it is essential to identify functional connectivity among retinal neurons. For this purpose, imaging neuronal activity through fluorescent indicator proteins has been a promising approach offering simultaneous measurements of neuronal activities from different regions of the circuit. In this study, we used genetically encoded voltage and calcium indicators, Bongwoori-R3 and GCaMP6f, to visualize membrane voltage and calcium dynamics in the form of the spatial map within retinal ganglion cells from retina tissues of the photoreceptor degenerated *rd1* mice. Retinal voltage imaging confirmed current-evoked responses from somatic spiking and intercellular conduction, while calcium imaging showed current evoked changes in calcium concentrations of presynaptic neurons. These results indicate that the combination of fluorescent protein sensors and high-speed imaging methods permits imaging electrical activity with cellular precision and millisecond resolution. Hence, we expect our method will provide a potent experimental platform for the study of retinal signaling pathways as well as the development of retinal stimulation strategies in visual prosthesis.

## INTRODUCTION

One of the most common retinal diseases is Retinitis Pigmentosa (RP) characterized by progressive degeneration of photoreceptors and sparing of retinal ganglion cells, which makes it the leading cause of legal blindness [1, 2]. One out of 4,000 people—a total of approximately one million individuals—are affected by RP [3]. Prosthetic implants including microelectrode arrays (MEAs) attempt to correct these irreversible diseases by using extracellular current to activate neurons in the visual pathways [4, 5]. In the blind retina, in which light-sensitive photoreceptors are damaged, electrical stimulation via retinal implants has been used to provide artificial visual input [6]. Retinal implants electrically stimulate the remaining retinal cells so that the brain interprets spiking activity of retinal ganglion cells (RGCs) into visual perception [7]. Among retinal prostheses, epiretinal implants are the most typical in which a microelectrode array is placed on the inner retina [8, 9]. The performance of these prostheses is evaluated by their ability to selectively activate populations of retinal ganglion cells with a high spatiotemporal resolution in order to faithfully convey visual information [10]. Retinal degeneration in the *rd1* mouse model of Retinitis Pigmentosa has been used to verify electrical stimulation onto retina tissue and investigate biophysical pathways of retinal signals [11, 12]. Understanding how retinal circuits transmit visual signals is crucial for developing vision restoration strategies.

As an electrical system, neuronal activity is most precisely measured by a pair of electrodes placed across the cellular membrane. However, the limited spatial resolution imposed by the physical dimensions of the electrodes has motivated the development of optical methods for the recording of physiological signals from neuronal populations. Instead of using electrophysiology, fluorescent protein-based indicators, which are sensitive to cellular calcium concentrations or membrane voltage, offer the possibility of imaging electrical signals from neurons [13]. Genetically-encoded voltage indicators (GEVIs) alter their fluorescent output in response to voltage transients at the plasma membrane, while genetically-encoded calcium indicators (GECIs) detect changes in intracellular calcium levels that generally increase in response to the firing of action potentials (APs) [14, 15]. Calcium sensors are commonly used to monitor neuronal activity, but do not directly match membrane voltage and cannot reliably report sub-threshold activities [16, 17]. Interpreting neuronal spiking activity can also be problematic because GECIs suffer from slow on-off kinetics [18], a shortcoming GEVIs can overcome to directly report neural activity [19-22]. Voltage sensors can more accurately follow membrane potential changes [13]. Earlier functional studies of the retina used the GECI, GCaMP3, to analyze retinal activity that was correlated across populations [23]. The improved GECI, GCaMP6s, was also used to observe retinal function [24]. More recently, the selective activation effect of retinal ganglion cells expected in electrical stimulation was optically tested with GCaMP6f [25]. In this paper, GCaMP6f, was used to compare with the voltage sensor, Bongwoori-R3.

A successful approach to visual restoration would be possible if we know the functional circuit of the retina and can physiologically activate them [7]. Therefore, it is important for developing retinal prosthetics to understand how retinal cell groups are activated in response to electrical stimulation. Still, there is an incomplete understanding of those functional connectivity in retinal neurons. Most studies on signaling circuits in retinal ganglion cells (RGCs) have used electrophysiology measurements [26, 27]. A previous study combining electrical imaging with extracellular multielectrode array [28] revealed the spatial propagation of electrical signals within the RGC [29]. However, since both stimulation and recording are electrical methods, the cross-talk occurs in which stimulus artifacts are mixed with desired signals from cells of interest [10]. This limits the simultaneous observation of retinal activity during the stimulation [10]. To overcome this issue, optical recording techniques have emerged based on fluorescent indicator proteins that can optically track cellular activities in real-time [30].

Recent advances in ArcLight-derived GEVIs have led to development of the GEVI, Bongwoori-R3, with better signal size, speed, and a voltage-response range tuned towards action potentials [31]. Imaging techniques utilizing these protein sensors have great potential in studying both single cell and population activity from retinal ganglion cells.

Here, we evaluated voltage and calcium indicator proteins as genetic tools for dissecting retinal connectivity in *rd1* blind retina. We utilized the advantages of the GEVI, Bongwoori-R3 to image retinal activity in response to electrical stimulation in the photoreceptor-degenerated mouse. To the best of our knowledge, the results presented here demonstrate the first attempt at voltage imaging using GEVI on RGCs. Retina voltage imaging directly visualized electrical activities within and across retinal ganglion cells, and showed that the optically recorded voltage signals from retinal ganglion cells correlated with their electrically recorded activities.

## RESULTS

### Imaging Activities from Retinal Ganglion Cells with Bongwoori-R3 and GCaMP6f

We expressed genetically encoded voltage and calcium indicators, Bongwoori-R3 (See also Supplementary Figure S1) and GCaMP6f, in retinal ganglion cells via viral injection, from *ex-vivo* retina tissues of the photoreceptor degenerated rd1 mice. To access the ability of these fluorescent indicators to optically report activities of retinal ganglion cells (RGC), RGCs expressing Bongwoori-R3 or GCaMP6f were observed under patch-clamp and one-photon fluorescence imaging (acquisition rate, 10 kHz and 1 kHz, respectively). A schematic representation of the experimental setup is shown in Figure 1.

**Figure 1.**
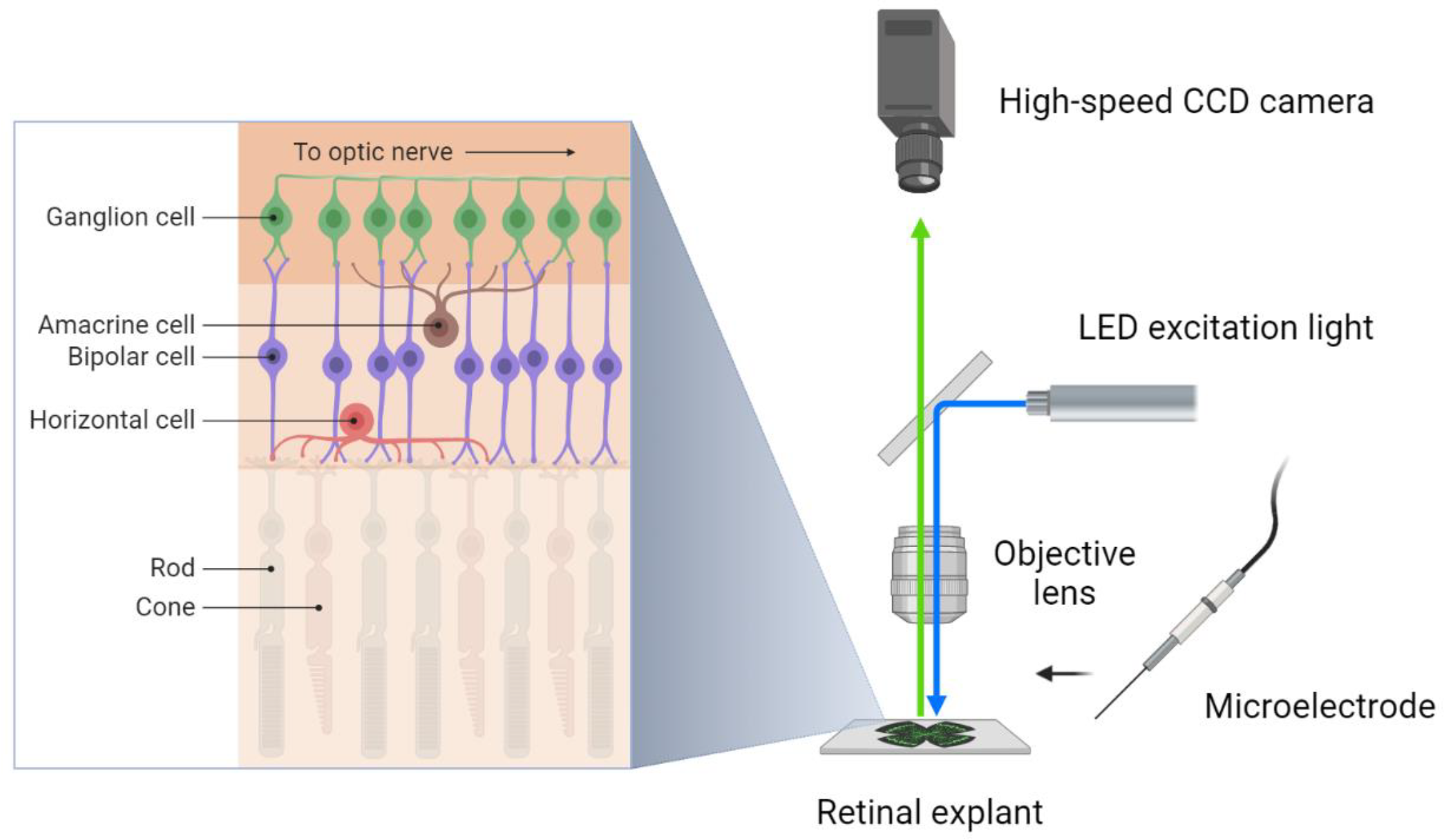
Schematic overview of the experimental setup. **(Left)** Visual information flows from photoreceptors to bipolar cells to ganglion cells and the brain. The retina consists of five layers of cells, including three nuclear layers and two synaptic layers. The nuclear layer has the photoreceptor, bipolar cell, ganglion cell, and the synaptic layer amacrine, horizontal cell. In the retina of blind retinal degeneration 1 mutation mice (*Pde6b*^*rd1*^ or *rd1*), rod and cone photoreceptors are degenerated. **(Right)** Retinal ganglion cell (RGC) of *rd1* mice was imaged during patch-clamp recording or extracellular electrical stimulation, with a bipolar microelectrode. A blue 460 nm LED was used for excitation of genetically-encoded fluorescent sensors virally expressed in a retinal explant. The emitted fluorescence from RGCs expressing either Bongwoori-R3 or GCaMP6f was recorded by a high-speed CCD camera.

### Both Bongwoori-R3 and GCaMP6f respond to action potentials

As shown in Figure 2, the electrophysiological traces show activities from the protein sensor expressing retinal ganglion cells. Mouse retinal ganglion cells expressing the GEVI, Bongwoori-R3 or the GECI, GCaMP6f were subjected to whole-cell current-clamp to image the fluorescence change of a single cell induced by the firing of action potentials (APs). In Figure 2A, each of the three trains of APs altered the fluorescence by about 1% ΔF/F. After each train of action potentials, the baseline is also slightly altered due to the acidification of the cell [31, 32]. Signals of both Bongwoori-R3 and GCaMP6f were found in the cell bodies of retinal ganglion cells. Also, Bongwoori-R3 signal was observed in their process, but GCaMP6f signal was less frequently detected (Figures 2A and 2C; Figures 4 and Supplementary Figure S2). These results reveal the spatial map in which the activity of retinal ganglion cell was optically tracked during a single trial (See also Supplementary Figure S3). While the calcium sensor exhibited slow dynamics with large fluorescence change, the voltage sensor exhibited faster kinetics, particularly at the end of electrical stimulus (See also Supplementary Figure S2).

**Figure 2.**
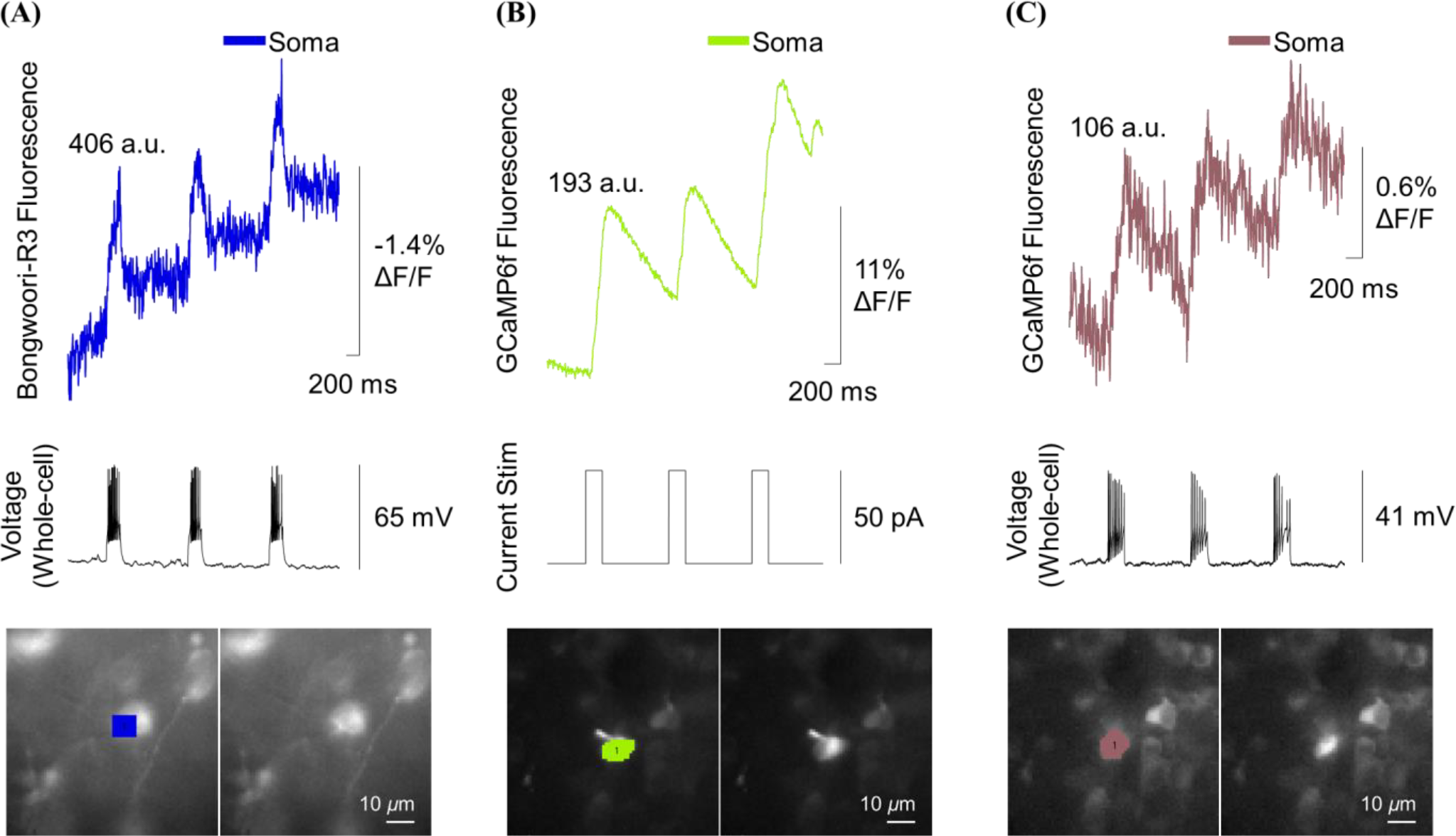
Electrophysiological and fluorescence signals from either the GEVI, Bongwoori-R3 or the GECI GCaMP6f expressing retinal ganglion cells of *rd1* blind mice in response to current-clamp steps. Retinal ganglion cells (RGCs) were imaged using a fluorescent protein called either (A) Bongwoori-R3 or (B, C) GCaMP6f at a rate of either 1 kHz. At the same time, the cell was also recorded using current-clamp mode at a rate of 10 kHz. Both signals were displayed in either color or black. The number above each trace shows the fluorescence intensity when the sample was not being stimulated. Both fluorescence and electrophysiology signals were obtained from each single trial. The bottom left image shows the specific pixels of interest. The bottom right image displays the resting fluorescence of retinal ganglion cells that express the Bongwoori-R3 protein. The fluorescence was measured by averaging the pixels of interest in the lower image. This is then displayed as color traces, which show how the fluorescence levels change over time. When the retinal ganglion cell was electrically stimulated, it resulted in the generation of fluorescence signals (color traces). The current stimulation was either (A, C) a strength of 100 pA for 200 ms in whole-cell patch, or (B) a strength of 50 pA for 200 ms in cell-attached patch mode. The calcium sensor responded more slowly to changes in electrical activity, while the voltage sensor responded more quickly.

The calcium signal showed a slightly larger and slower optical response when retinal ganglion cells expressing GCaMP6f were imaged under a cell-attached configuration (Figure 2B) and whole-cell current-clamp conditions (Figure 2C). There is a slight technical challenge for the calcium imaging. The negative pressure required to obtain a tight seal between the electrode and the cell would result in a washout of the GECI upon going whole-cell. This did not affect GEVI recordings as that probe resides in the plasma membrane. Fortunately, action potentials could be induced in the cell-attached configuration resulting in robust calcium signals of greater than 10% ΔF/F (Figure 2B). GCaMP6f signal could also be seen in the whole-cell configuration despite the significant reduction in the signal size (Figure 2C).

Data from Figures 2A and 2C were used to make Supplementary Movies S1 and S2, respectively (See also Figure 10, A and B). The images were acquired at 1 kHz and binned every 10 frames to create the movies. These movies show fluorescent voltage and calcium indicators reporting the entire process leading to membrane potential transients (See also Figures 10A and 10B). In all the movies, electrophysiological recording (bottom) and one-photon fluorescence imaging (top) are concurrently provided at each time point (marked by either red or white square in the bottom trace).

### Imaging sub-and supra-threshold activity as well as intercellular communication in retinal ganglion cells with Bongwoori-R3

Bongwoori-R3 can optically report isolated action potentials of retinal ganglion cells (Figure 3B, first trace of purple). However, resolving multiple action potentials (APs) at high frequency is a challenge due to the reduction in the signal-to-noise ratio (Figure 2A; See also Supplementary Figure S3). Unlike the cultured hippocampal response where the investigator can choose a neuron that is well isolated from other fluorescent cells with low internal fluorescence, the architecture of the *ex-vivo* retinal preparation increases the background noise in the recording. Background fluorescence can affect the signal and noise levels. An example of this can be seen in Figure 3B where only 11 pixels give a reasonable optical representation of the voltage transients in the patched retinal ganglion cell experienced. Adding pixels to the region of interest reduces the noise in the baseline of the trace by increasing the number of photons, but also alters the signal size of the spiking activity.

**Figure 3.**
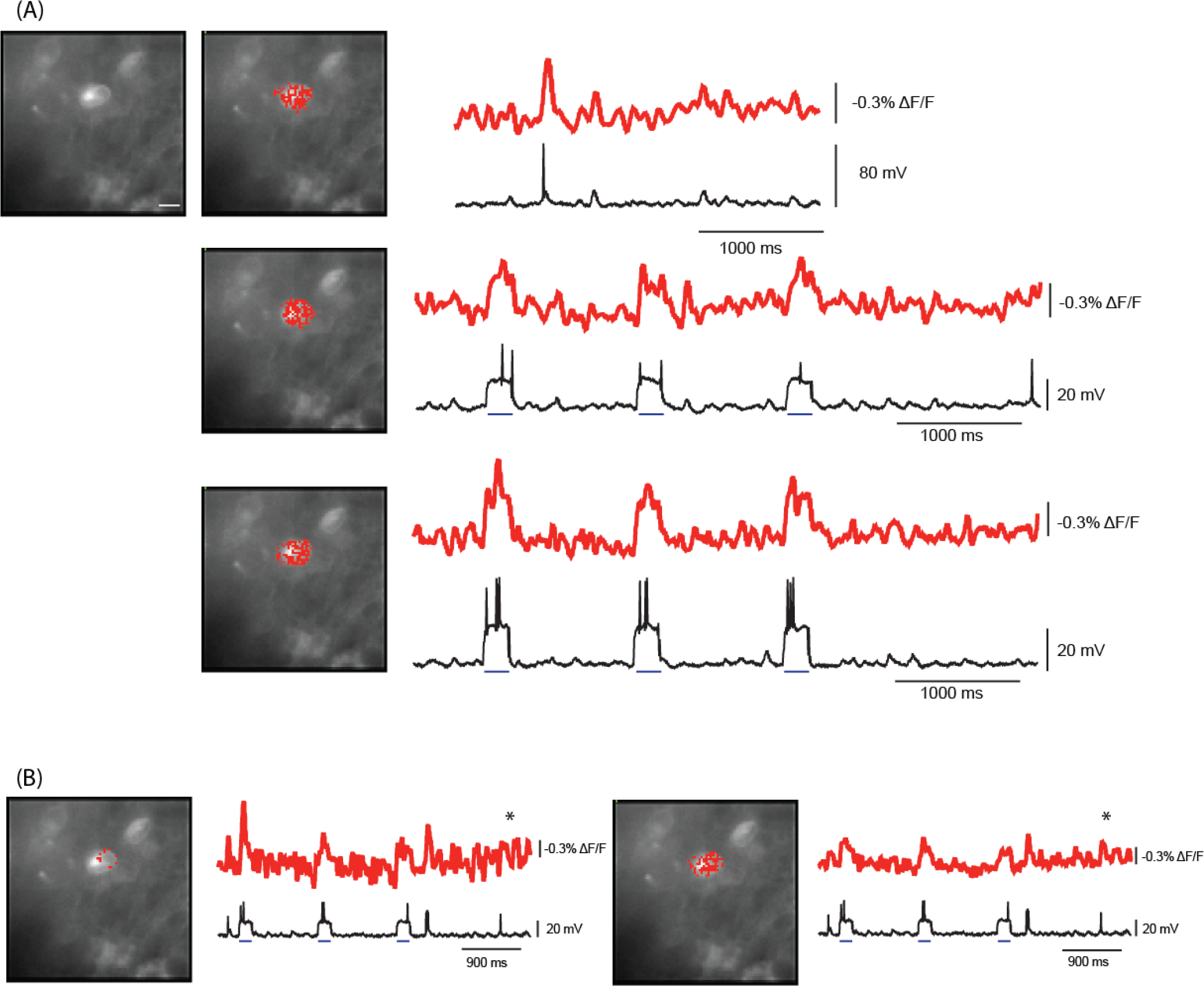
Imaging voltage with the GEVI, Bongwoori-R3 expressing retinal ganglion cells of *rd1* blind mouse. (A)Spontaneous (top trace) or induced activity (lower two traces) were imaged at 1 kHz See also Supplementary Figure S3. The region of interest (red pixels) correlates to the retinal ganglion cell expressing Bongwoori-R3 under whole cell current clamp configuration. Injected current (blue bar) induced the firing of action potentials. (B). The region of interest influences the signal to noise ration. The same trials is shown twice. The left trace is from 11 pixels yielding the largest optical change for the first current injection. The right trace includes more pixels to reduce the baseline noise in an attempt to resolve the last spontaneous voltage transient (asterisk). Scale bar is 10 μm.

Despite these challenges, Bongwoori-R3 is able to differentiate sub-threshold activity from APs as well as report intercellular communication. Figure 4A and 4C shows another population of retinal ganglion cells expressing Bongwoori-R3. The cell in the center was subjected to whole-cell current-clamp and given sub-threshold stimulation (Figure 4B, black trace at the bottom) that could only be optically detected in the soma (Figure 4B, ROI 1). Due to the small voltage change of only around 20 mV (Figure 4B), ‘frame subtraction’— mean fluorescence of the frames during the first stimulation were subtracted from the mean fluorescence of frames having no observable electrical activity—was used to identify pixels responding to the membrane potential transient. The resulting pixels corresponded primarily to the plasma membrane. Increasing the stimulation to the patched cell resulted in the firing of action potentials (Figure 4D). The corresponding optical signals to the stronger stimulation could be detected in the axon as well two regions outside the patched cell (Figure 4D, ROIs 1–4).

**Figure 4.**
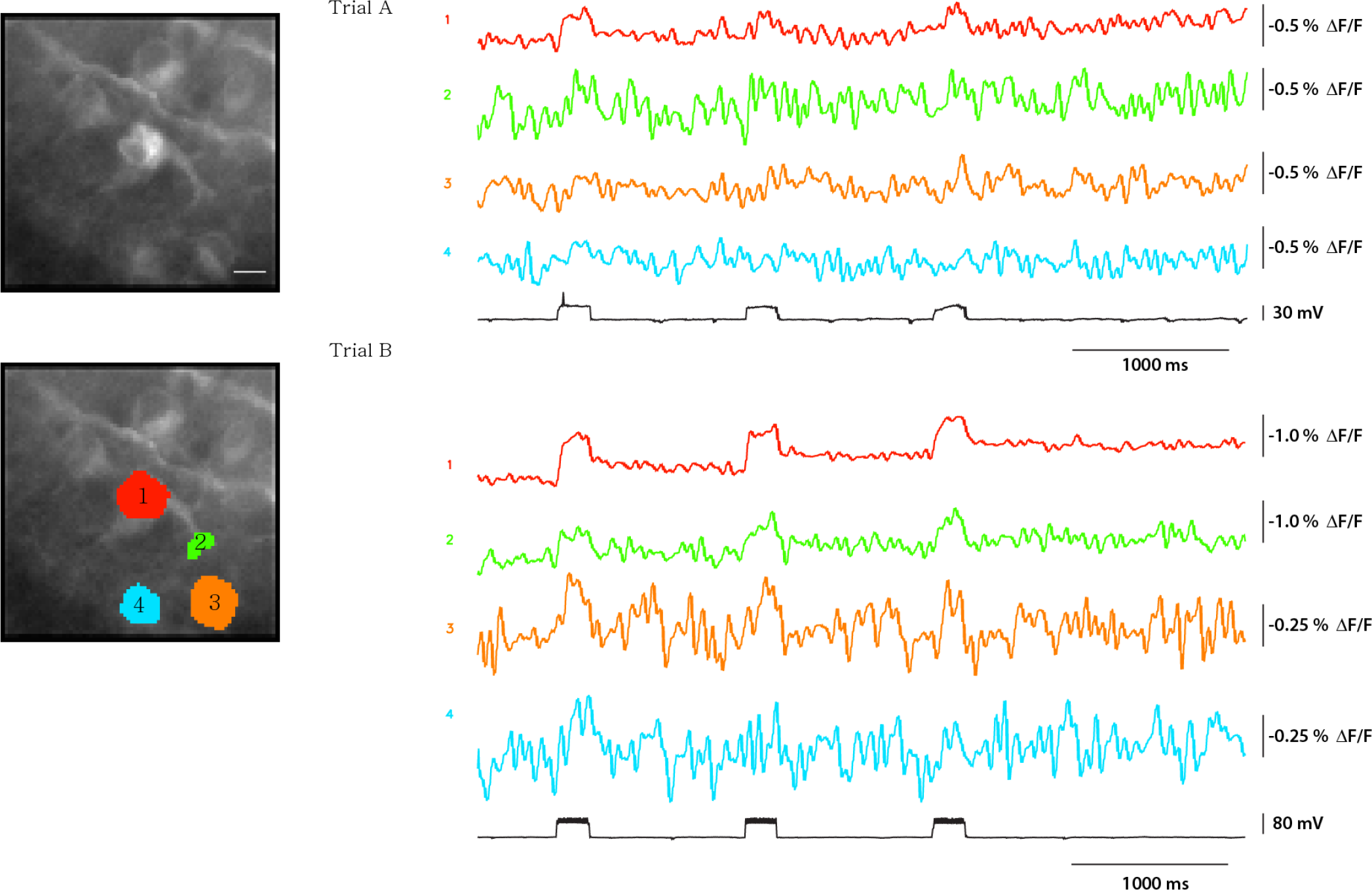
Tracing downstream signals from the soma with Bongwoori-R. Retinal ganglion cells (RGCs) expressing Bongwoori-R3 were imaged at 1 kHz. The cell corresponding to ROI 1 (red pixels) was under whole current clamp. Trial A injection of current yielded only one action potential. The voltage transients in trial A can only be detected in the soma of the patched cell. In trial B a stronger stimulation elicited multiple action potentials. Voltage transients can be detected in the soma (red trace), the axon (green trace) and a neighboring cell (orange trace). Another neighboring cell (blue trace) may show a signal during the first pulse but not the second or third lulses. All traces were subjected to an offline low pass butterworth filter (20Hz). Scale bar is 10 μm.

### Identifying gap junctions with Bongwoori-R3

In addition to tracing the retinal activity from the soma to the axon to neighboring cells, Bongwoori-R3 could also identify the presence of at least one gap junction between non-neighboring RGCs. Prior work on gap junctions in RGCs [33] found that electrical synapses are responsible for the concerted spiking between RGC neighbors in the retina. In Figure 6, whole-cell voltage-clamp was employed to manipulate the membrane potential of a single RGC. When the Soma 1 (pink trace) is subjected to voltage steps, an optical signal can be seen in the distal Soma 4 (blue trace) as well as in the projection extending from that cell. Neighboring soma do not respond (Soma 3; magenta trace) though some light scattering can be seen in one neighboring soma (Soma 2; green trace). Also, the light scattered from the fluorescent signal in Soma 5 is seen on the adjacent cell body. Soma 4 in the upper left quadrant in the bottom of Figure 6A exhibits a fluorescence response when the membrane potential of the Soma 1 is hyperpolarized as well as during depolarization steps (Figure 6B, black trace at the bottom) indicating the presence of an electrical synapse between retinal ganglion cells between it and the patched retinal ganglion cell.

### Identifying axonal projections with Bongwoori-R3

Retinal ganglion cells bundle their axons to become the optic nerve. Since Bongwoori-R3 was able to report voltage transients in the axon (Figure 4D, ROI 2), we employed whole-cell voltage-clamp in order to determine the extent we could affect a fluorescence response into axon bundles. Figure 7A shows a population of retinal ganglion with two tracts of fluorescence corresponding to distinct axon bundles. The trace in figure 7B is an average of 16 trials. The cell under whole-cell voltage-clamp (ROI 1; red trace) projects its axon in the lower bundle and not the upper bundle (compare ROI 3 to ROI 4; violet and light blue traces). This is a significant technical achievement as the bundle consists of multiple axons from multiple cells and only one of which is experiencing a voltage change. Despite nonresponsive fluorescence of the other axons in the bundle, a clear signal can be seen in the Figure 7B (ROI 3; magenta trace) when the Soma 1 (red trace) is subjected to voltage steps (black trace). Indeed, we were even able to see the hyperpolarizing steps in certain sub-regions of the axon bundle (Supplemental Figure S4; ROI 5).

### Differentiating the voltage-dependent and pH-dependent optical signals with Bongwoori-R3

There are different approaches for determining the regions of interest for optical interrogation. One approach is to use anatomical features such as soma of distinct cells (Figures 2–3, 5) or the axon bundles seen in Figures 4 and 6. However, optical imaging also offers the use of ‘frame subtraction’ to identify regions of fluorescence change. By averaging the fluorescence from a collection of frames preceding or prior to a stimulus, one can then subtract the fluorescent average during the stimulus and plot the difference to identify the pixels detecting a change in the fluorescence. Regions that have no change in fluorescence subtract to zero.

**Figure 5.**
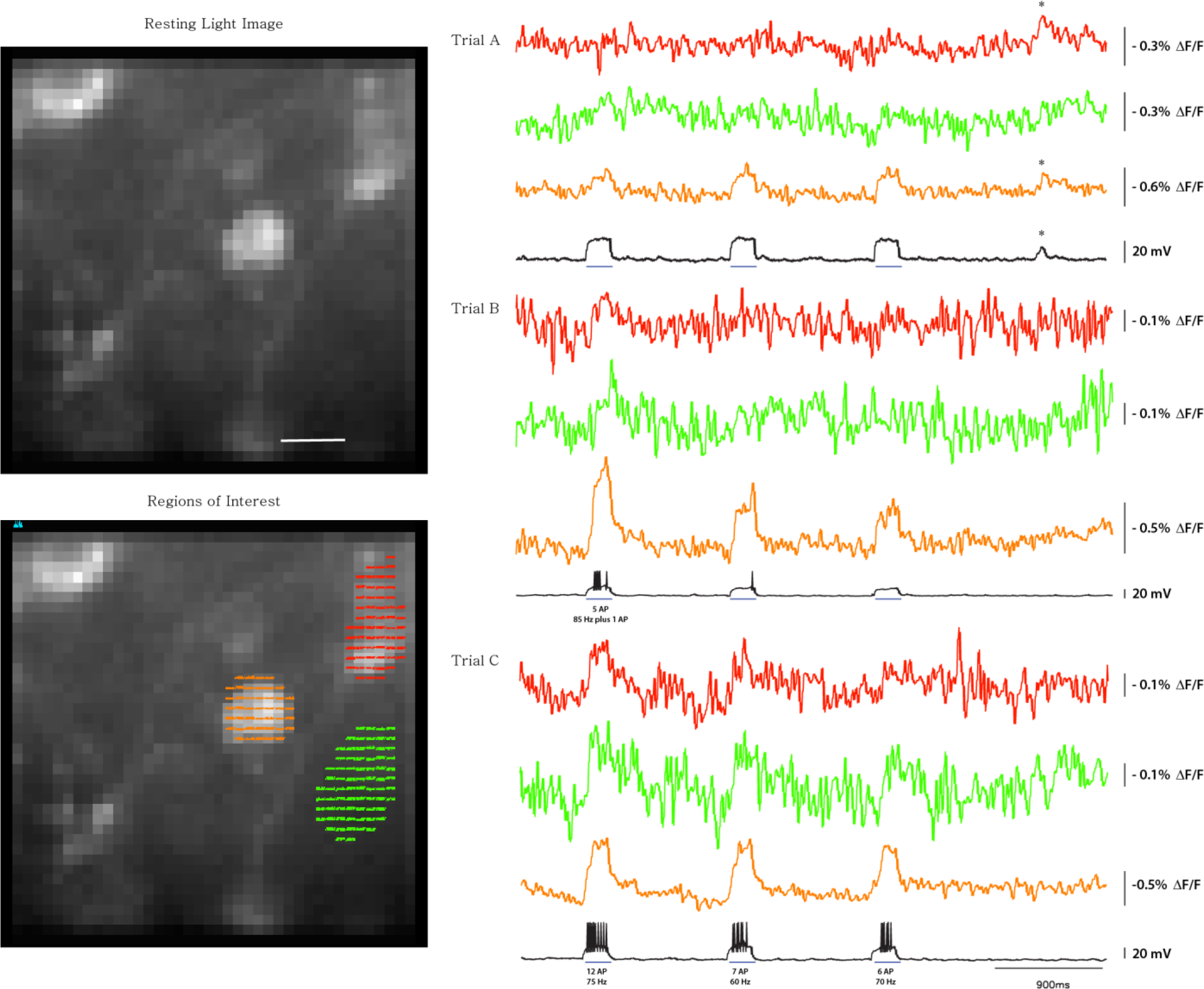
Intercellular communication in the retina of *rd1* blind mice. Retinal ganglion cells (RGCs) expressing Bongwoori-R3 were imaged with a frame rate of 3 kHz (same cells as in figure 2A). The center cell (orange ROI and corresponding traces) was under whole cell current clamp configuration. Current injection (blue bars) was varied each trial resulting in subthreshold only voltage transients in Trial A and action potentials (AP) with stronger stimulations in Trials B and C. In Trial A, the voltage transients could only be optically detected in the patched cell (orange trace) with the exception of a spontaneous event (asterisks) that was detected in the red region of interest as well. In Trial B, the stimulation was close to the threshold level resulting in multiple action potentials during the first injection, one action potential during the second injection, and only a subthreshold change in the third. Trial C yielded multiple action potentials during all three current injections. The faint cell in the green region of interest was able to report voltage transients only when multiple Aps were elicited in the patched cell. The red region of interest also reported voltage transients when the patched cell fired multiple action potentials except for the third pulse in Trial C which may be due to its higher internal fluores-cence lowering the SNR. Other regions in this field of view also yielded optical signals (See Supplementary Figure S3). Voltage re-cording is in black. Optical traces were subjected to an offline low pass Butterworth filter (30 Hz) as well as an exponential subtrac-tion to correct for baseline drift. For 3 kHz recording the pixels are binned resulting in a 40x40 pixel image as compared to an 80x80 pixel image for the 1 kHz recordings. The white bar in the image is 10 μm.

Using ‘frame subtraction’ enables the discernment of a voltage-dependent fluorescence change from a pH-dependent fluorescence change (Figure 8). Bongwoori-R3 is an ArcLight-derived GEVI consisting of a transmembrane domain that responds to voltage transients linked to a cytosolic fluorescent protein domain that is pH sensitive. Bongwoori-R3 therefore has a pH-sensitive component as well that does not hinder its ability to respond to voltage [34] but will change the fluorescence baseline if the cell experiences a change in pH [31].

In Figure 8A, the average fluorescence of 100 frames during a voltage step (Figure 8B, orange bar labeled b) was obtained by subtracting the average fluorescence of 100 frames before or after a voltage step (Figure 8B, blue or green bar labeled a or c), as shown in the middle or bottom image. In the bottom trace of Figure 8B, plotting the difference in fluorescence of the frames during the depolarization of the plasma membrane (orange bar labeled b) from the frames preceding the voltage step (blue bar labeled a) reveals an optical signal in the soma of the retinal ganglion cell under voltage clamp (Figure 6B, Soma 1) as well as a signal in the upper left corner (Figure 6A, Soma 4) resulting from the gap junction. Based solely on this figure, one would most likely create a region of interest consisting of the entire soma and the process. However, if the frames after the voltage step are used (Figure 8B; green bar labeled c), the optical signal is different. This time the optical signal is limited to the plasma membrane while the cytoplasm does not exhibit much fluorescence change upon repolarization of the cell to the holding potential. This is due to the acidification of the cell during the depolarization step. By creating two regions of interest (Figure 8A, top), the green pixels corresponding to the plasma membrane and the red pixels to the cytoplasm, one can see that the kinetics of the optical signal are different (Figure 8B, first and second traces). For the signal from the plasma membrane (green pixels) the change is relatively fast and recovers more upon repolarization of the cell (Figure 8B, first trace). The cytoplasm, in contrast, acidifies resulting in a slower fluorescence change that persists (Figure 8B, second trace). The initial change in fluorescence is the combination of the change in membrane potential and the change in pH (ΔF/F = ΔF/F(pH) + ΔF/F(V)). The change in fluorescence during the repolarization to the holding potential is primarily voltage (ΔF/F = ΔF/F(V)). The change in baseline is then due to the persistent acidification of the cell during the length of the recording (ΔF/F = ΔF/F(pH)).

**Figure 6.**
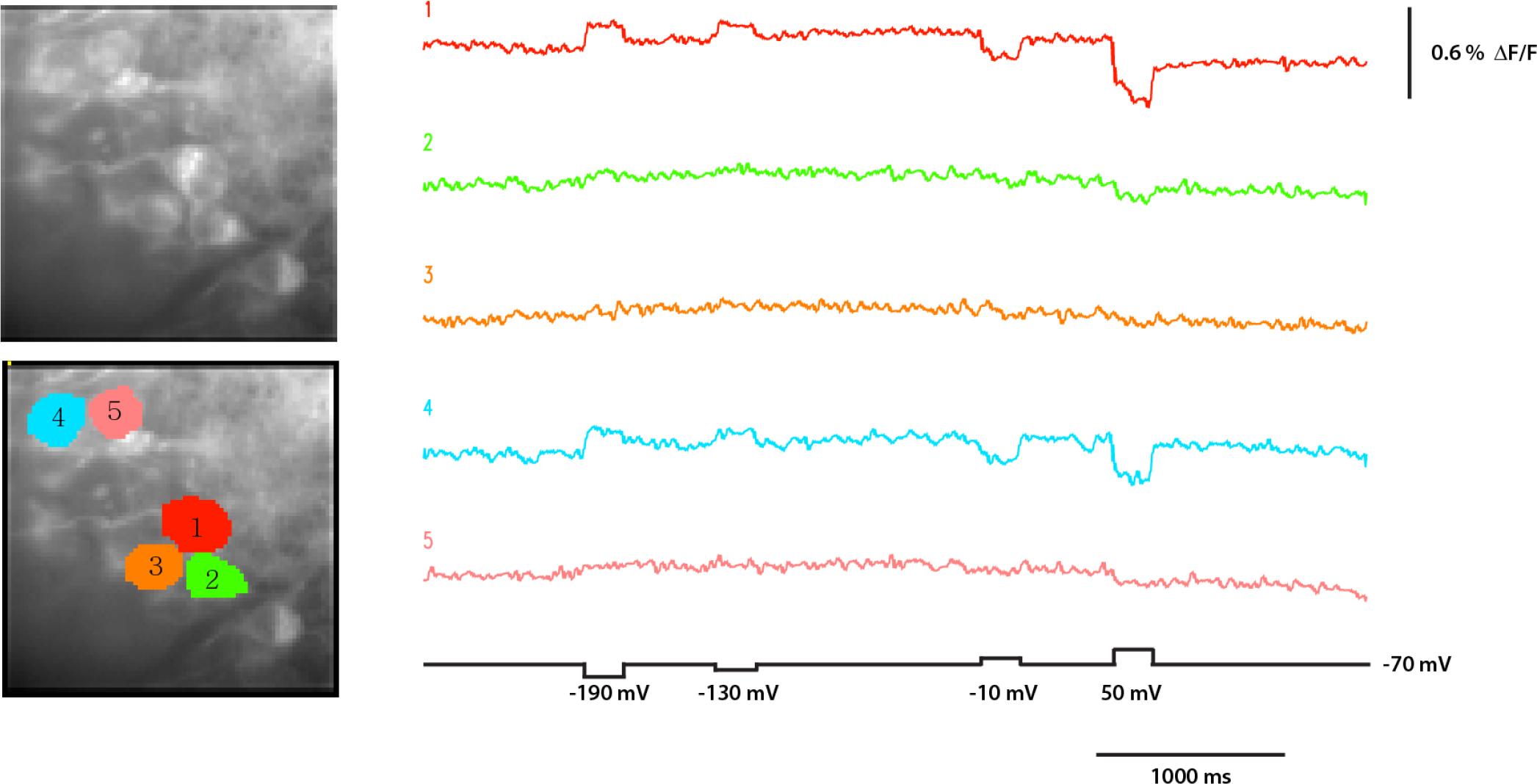
Imaging an electrical synapse between RGCs. The retinal ganglion cell in the red region of interest was subjected to whole cell voltage clamp The fluorescence traces are the average of 16 trials (frame rate 1 kHz) with an offline low pass Butterworth filter (20 Hz). Voltage steps are as indicated in the black. Cells 1 and 4 are clearly electrically coupled while the slight optical signal for the 120 mV depolarization step of the plasma membrane in cells 2 and 5 are likely due to light scattering from cells 1 and 4.

### Field stimulation activates multiple fibers improving the signal size of the optical response

Depending on the nature of the investigation, one may want to excite multiple cells to observe the response of the network. Bongwoori-R3 enables the detection of voltage transients in retinal explant under electrical stimulation with epi-retinal microelectrode (Figure 9) (See also Supplementary Figure S5). The blue region in Figure 9A corresponds to the blue traces in Figure 9C. The effect of stimulus phase and frequency on retinal cells was analyzed by measuring changes in fluorescence intensity, which is indicative of membrane potential changes. The minimum current stimulation to evoke fluorescent voltage signals was 0.5 ms long with an amplitude of 2 mA. These were the conditions for our particular sample. Figure 9C shows evoked voltage signals under pulse stimuli (2 mA; a threshold for depolarization), with the pulse rate of 1 Hz in the left panel and the pulse rate of 10 Hz in the right panel. In each panel, the first, the second, and the third columns display voltage imaging signals observed with pulse durations of 1 ms, 5 ms and 10 ms. The first and the third rows show voltage imaging signals from positive and negative phased current pulses, respectively. For the single-trial traces in Figure 9C, the fall time was quantified as time intervals from the optical peak to baseline signal. In addition, decaying fractional value (ΔF/F) was quantified as fluorescence difference between the first and the last 100 frames. These optical characteristics from one-photon microscopy were analyzed and summarized in Figure 9B. These results show that Bongwoori-R3 optically reports voltage responses to electrical stimuli during a single trial. Consistent with the prior studies, fluorescence baseline decreased (Figure 9C) due to cellular acidification during firing of action potentials [31, 35].

In Figure 9C, retinal activity indicated by fluorescence showed 9, 26, and 17 ms of peak delayed after 1 Hz cathodal pulses of 1, 5, and 10 ms, respectively, and 8, 12, and 13 ms of peak delayed after 1 Hz anodal pulse of 1, 5, and 10 ms, respectively (both; not shown). Also, the fluorescence signal representing retinal activity had delayed peaks of 11, 11, and 21 ms after 10 Hz cathodal pulses of 1, 5, and 10 ms, respectively, and delayed peaks of 11, 13, and 14 ms after 10 Hz anodal pulses of 1, 5, and 10 ms, respectively (both; not shown). The delay time of 1 Hz anodal pulses was significantly faster than 1 Hz cathodal pulses (*p* = .0391). The delay for 1 ms pulses of 1 Hz was significantly shorter than that for 5 ms pulses (*p* = .0197) and also significantly shorter than that for 10 ms pulses (*p* = .0002). The delay for 10 ms pulses of 10 Hz was significantly longer than that for 1 ms pulses (*p* = .0188) and also significantly longer than that for 5 ms pulses (*p* = .0450).

Imaging data with epi-retinal stimulation (Figure 9C; cathodal and anodal pulses; amplitude, 2 mA; duration, 10 ms; rate, 10 Hz) were used to make Supplementary Movies 3 and 4. Data from Figures 10C and 10D were used to generate Movies S3 and S4, respectively (See also Figure 10, B and C). The images were acquired at 1 kHz and binned every 10 frames to make the movies. These movies visualize the spread of network activity in the retina detected with the fluorescent voltage-indicator protein during an electrical stimulation (See also Figures 10C and 10D).

As displayed in the Movies S3 and S4, epi-retinal stimulus waveform (bottom) and one-photon fluorescence imaging (top) are simultaneously presented at each time point (marked by a red square in the bottom trace). The stimulating microelectrode was placed above the retina tissue in an epi-retinal configuration and is located outside the field of view (FOV) at the bottom right, the direction marked as STIM in the movie clips. Regardless of the pulse phases, electrical stimulation generated retinal responses involving retinal cells and axon bundles. Cathodal pulses induced consistent responses compared to anodal pulses. This result demonstrates that GEVIs can optically track network activity in a spatially defined manner within the retina in response to electrical stimulation.

## DISCUSSION

Visual prosthetic devices activate retinal cells as a part of artificial visual information for restoring visual perception. Improving the performance of these implants requires an understanding of the operational principles of the retinal circuit dynamics, each with individual visual processes. For this purpose, genetically-encoded protein sensors were utilized in this study to spatially map retinal population activity: GECIs were used to detect calcium changes which exhibits slow dynamics associated with neuronal cell activity, and GEVIs were used to detect fast voltage changes as direct evidence of cell firing (Figures 2 to 4). This study reveals that voltage and calcium one-photon fluorescence imaging are suitable for retinal explants from photoreceptor degeneration mice. These findings have important implications: cellular resolution, and cellular connectivity detectability (Figures 2 to 7). Furthermore, *rd1* blind mouse, the murine model of retinal disease [1] permits an investigation of retinal output neurons without the contaminating activation of photoreceptors [36]. Moreover, these approaches overcome electrical artifacts during stimulation and recording (Figure 9). Thus, the results demonstrated in this work should encourage further investigation into retinal circuit function using genetically-encoded protein sensors.

**Figure 7.**
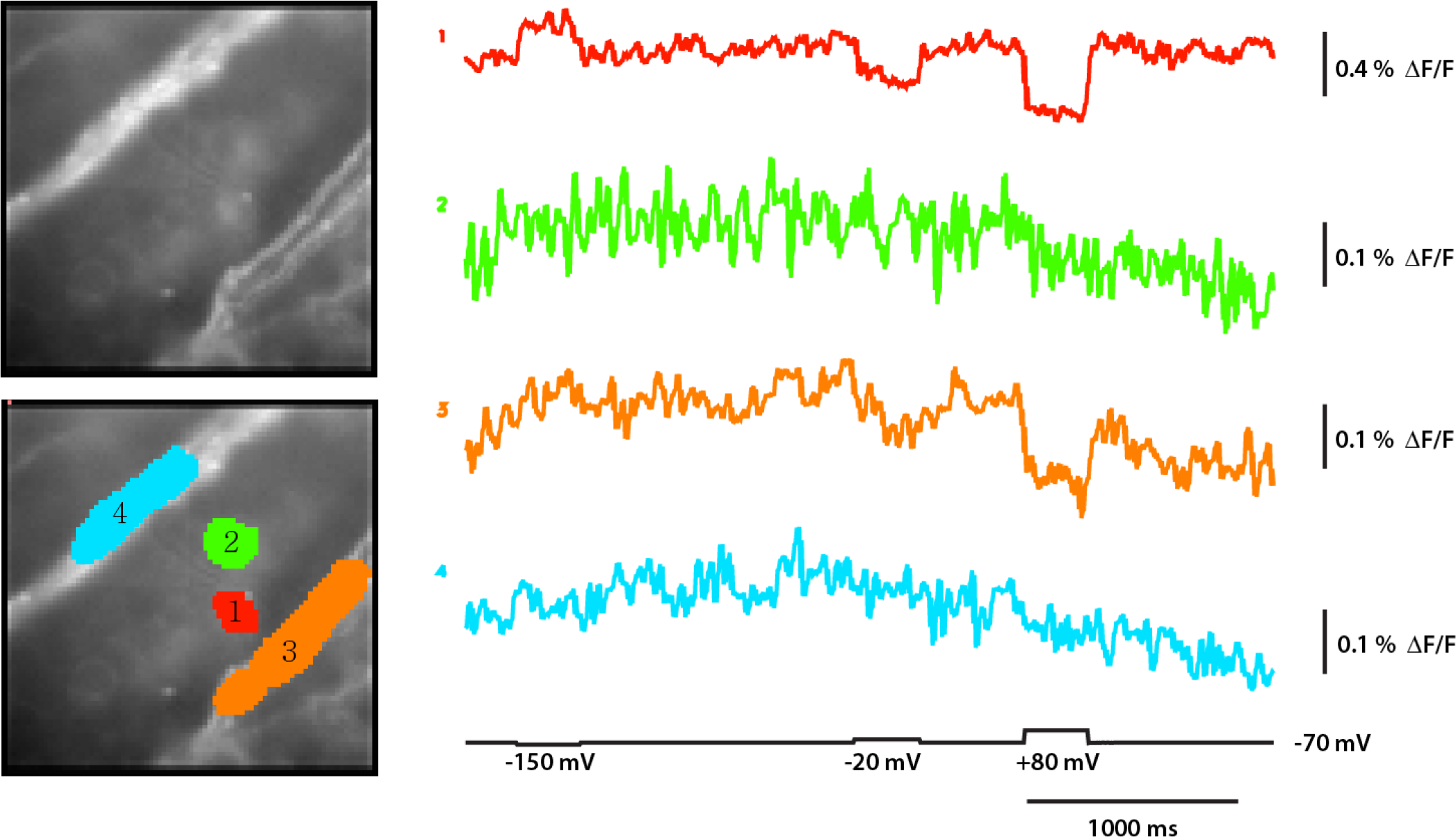
Fluorescence signals from Bongwoori-R3 expressing retinal ganglion cells and axon bundles of *rd1* blind mice in response to voltage-clamp steps. Retinal ganglion cells (RGCs) were imaged using a fluorescent protein called Bongwoori-R3 and simultaneously recorded using voltage-clamp. The fluorescence data was averaged from 16 trials, each sampled at 1 kHz. The data was then filtered using a Gaussian filter with a low pass set at 20 Hz. The fluorescence signal is presented in color. **(A)** The upper image shows the resting fluorescence of retina expressing Bongwoori-R3, while the lower image shows specific pixels of interest corresponding to RGCs and axon bundles. **(B)** The fluorescence was obtained by averaging the pixels of interest. This is then shown as color traces. The voltage of Soma 1 (red trace) was controlled to follow the step protocol at the bottom (black trace), which had an amplitude of -50, 50, and 150 mV, a duration of 300 ms, and a holding potential of -70 mV. Applying voltage steps to Soma 1 (red trace) caused a fluorescence signal to appear in Axon bundle 1 (magenta trace). This indicates that the cell’s axon selectively projects to the lower axon bundle, not the upper bundle (ROI 3–4; magenta and green traces). For further refinement of the optical signal in the axon bundle see Supplementary Figure S4.

This set of experiments present characteristics of two protein-based indicators, the GEVI, Bongwoori-R3 and the GECI, GCaMP6f. The results of the experiments indicate that the selection of an indicator sensor is important in monitoring cellular activities. Important aspects of this study include the ability of the GEVI, Bongwoori-R3 to trace axonal projections (Figures 4 and 6) and identify electrical synapses (Figure 6) as well as the response to action potentials (Figures 2 to 4).

As mentioned by another group [37], optical signals from explanted tissues, in general, exhibit a decrease in brightness and signal size because of tissue auto-fluorescence compared to cultured neurons. There is also an intracellular fluorescence that yields a small but noticeable pH response (Figure 8). Despite these limitations, the work presented here shows that Bongwoori-R3 worked well at the tissue level as well as in the presence of axonal bundles (Figure 7). In retinal explants, voltage imaging signals from Bongwoori-R3 yielded ΔF/F peak of ∼13% (Figure 9C) including pH-dependent fluorescence change of ∼3.8% (Figure 9B; decay ΔF/F), under epi-retinal electrical stimulation.

**Figure 8.**
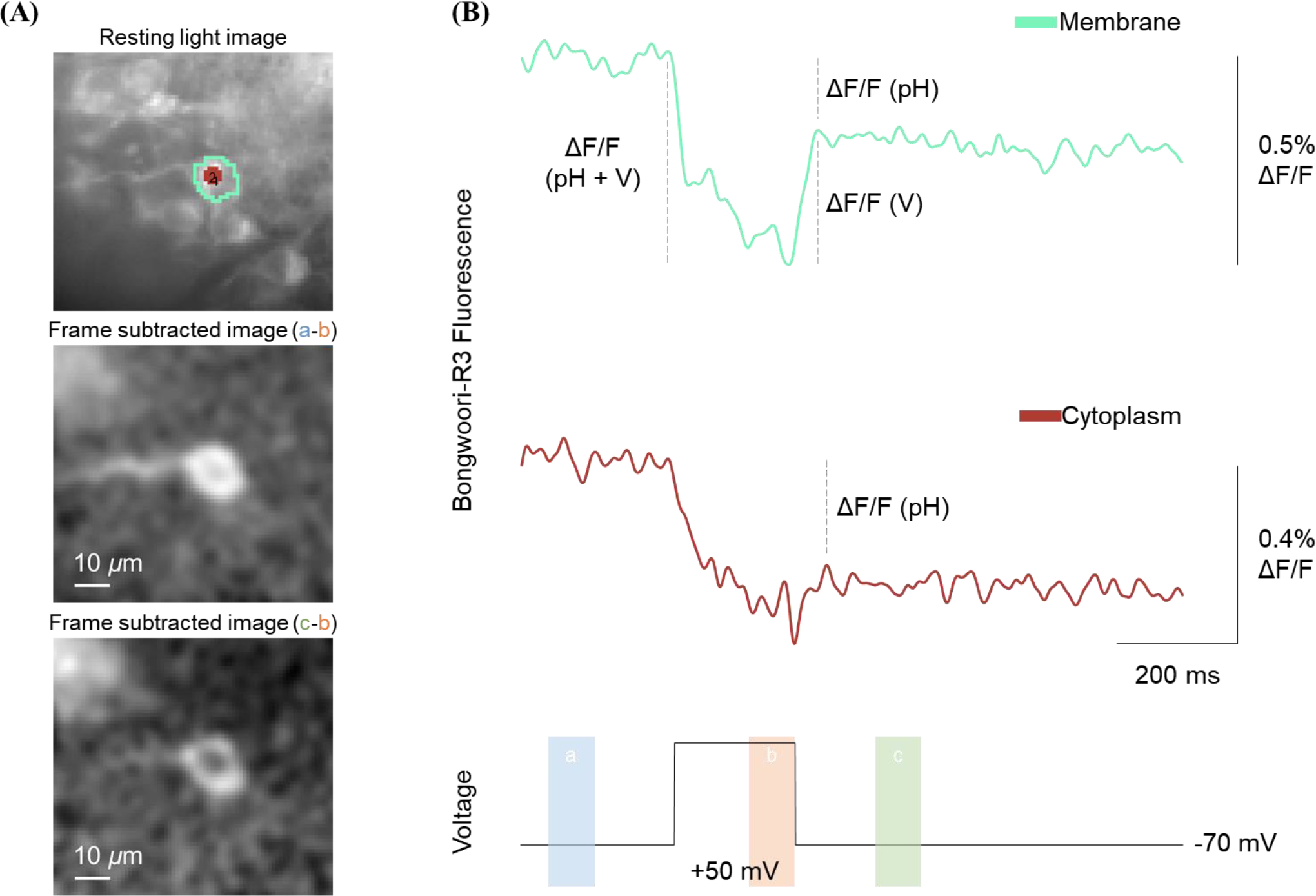
Observing physiological changes in a retinal ganglion cell in *rd1* blind mice: discerning voltage induced fluorescence change from pH. Fluorescence and electrophysiology signals were acquired at the same time. The fluorescence trace from 16 trials was collected at 1 kHz, averaged, and then filtered using a Gaussian filter with a low pass cutoff of 20 Hz. The fluorescence signal is shown in color. This figure was produced by the same data as in Figure 6. **(A)** The top image shows the resting fluorescence of RGCs that express Bongwoori-R3 protein. In the image, the pixels of interest correspond to the membranes and cytoplasm of the RGC. The middle and bottom images show the fluorescence intensity difference either between bars b&a, or between bars b&c depicted in the voltage traces in Figure 8B. **(B)** The fluorescence signal was obtained by averaging the pixels of interest. This is then displayed as color traces. The voltage of Soma 1 (Figure 6B; pink trace) was manipulated to match the step protocol at the bottom (black trace), which had the following parameters: amplitude of 120 mV; duration of 200 ms; holding potential of -70 mV. Comparing frame subtraction images reveals where in the RGC fluorescence changes occur before or after a voltage step (Figure 8A; middle and bottom images). The fluorescence change was mainly observed at the RGC’s plasma membrane in one image (Figure 8A, bottom image), created using frames after the voltage step (Figure 8B, green bar labeled c at bottom). On the other hand, the pH fluorescence response was seen in the RGC’s cytoplasm compared to its plasma membrane in another image (Figure 8A, middle image), created using frames before the voltage step (Figure 8B, blue bar labeled a at bottom). When returning to the holding potential, larger voltage responses were detected at the plasma membrane than in the cytoplasm (Figure 8B; ΔF/F (V) of Membrane and Cytoplasm).

The latency before the first spike of *rd10* RGCs was investigated with respect to the stimulation polarity by Stutzki et al. [7]. In that paper, an anodal stimulus of 2 ms duration caused a response delay of 7 ms, and a cathodal caused a response delay of 35 ms. They explain that if the stimulation phase is cathodic, epiretinal stimulation instead stimulates bipolar cells or adjacent interneurons to activate a small number of RGCs, eventually showing long-delayed spikes. In contrast, it has been reported that RGCs can be directly stimulated by applying a cathodic pulse using an epiretinal electrode [38]. Nevertheless, the response of a short delay was considered to have directly stimulated ganglion cells [39]. In this regard, in Figure 9C, there was a millisecond delayed peak for both positive and negative stimuli, which may have activated cells in the inner nuclear layer (INL) under the ganglion cell layer (GCL) along with the layered structure.

**Figure 9.**
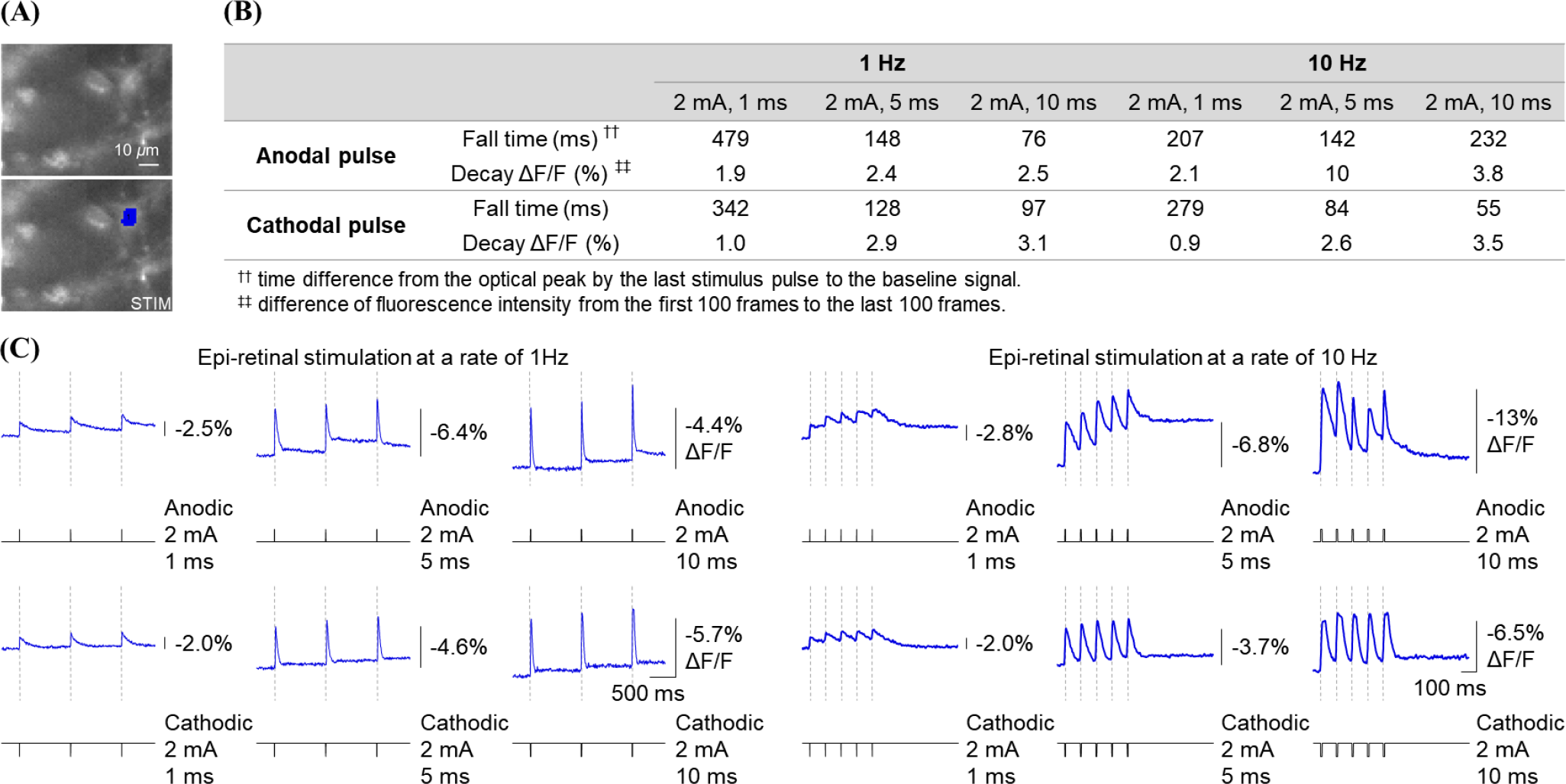
Fluorescence signals from Bongwoori-R3 of retina tissue in *rd1* blind mice during electrical stimulation with epiretinally positioned microelectrode. Retinal ganglion cells (RGCs) were imaged using a fluorescent protein called Bongwoori-R3 and concurrently stimulated using microelectrode. Fluorescence trace was recorded from a single trial, each sample taken at a rate of 1 kHz. The signal is displayed in color. **(A)** The upper image shows the resting fluorescence of retina expressing Bongwoori-R3, while the lower image shows the specific pixels that were used for analysis. A stimulating microelectrode was placed in the lower right corner of the image, outside the field of view. This is indicated by the “STIM” label. The fluorescence was measured by averaging the pixels of interest in the image. This is then displayed as a color trace, which shows how the fluorescence level change over time. **(B)** How different types of current stimulation affected the retina’s fluorescence response was analyzed, with a focus on the last peak in the fluorescence response. **(C)** When the retina was electrically stimulated using an applied current (black trace), it responded by generating fluorescence signals (color trace). The stimulus was described on the right side of each trace (amplitude, 2 mA; duration, 1, 5, 10 ms; rate, 1, 10 Hz). For reference, the vertical dash lines mark the start of the stimulus pulse. How retinal cells respond to different stimulus frequencies and parameters was compared by measuring their fluorescence response. This fluorescence signal indicates changes in the membrane potential of the cells.

**Figure 10.**
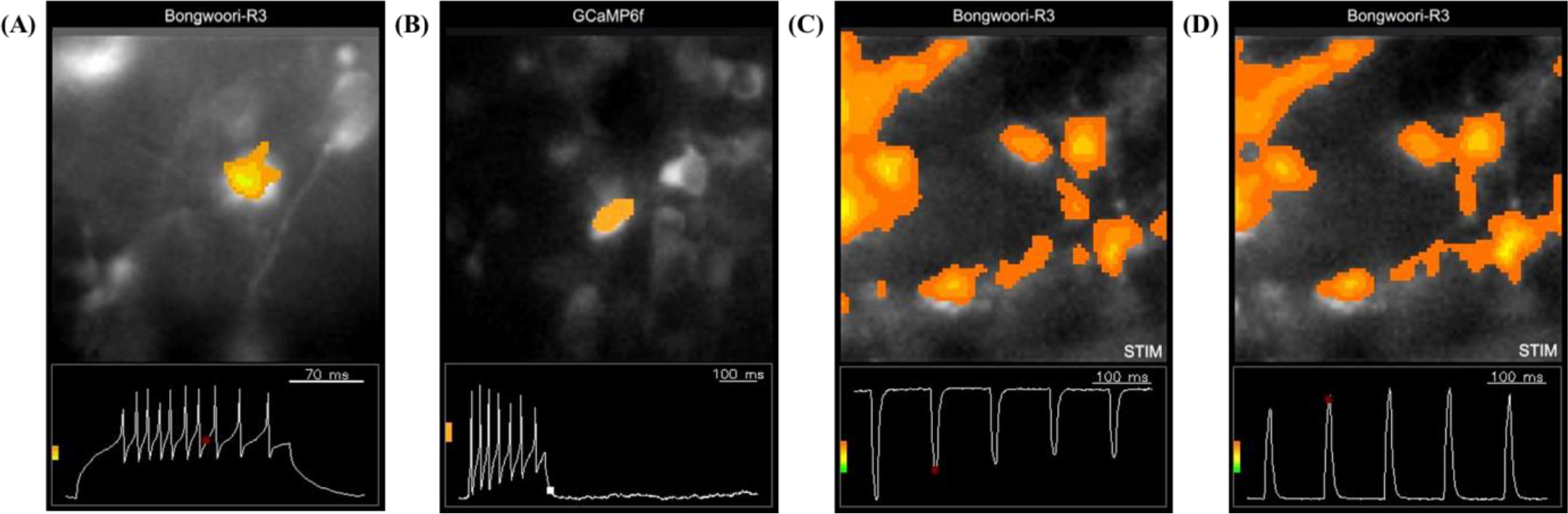
Imaging action potentials from either Bongwoori-R3 or GCaMP6f expressing retinal ganglion cells of *rd1* blind mice in response to electrical stimulation with current-clamp steps or epi-retinally positioned microelectrode. Imaging frames A to D were used to create Movies S1 to S4. The location of fluorescence changes was seen over time along the action potential (Figure 10, A and B) or stimulus (Figure 10, C and D), indicated by the small red or white box in the lower section. To remove noise and focus on the stimulated cells and their surrounding axon bundles, the threshold for fluorescence change was adjusted. This is represented by the color bars on the left. These figures were produced using the same data as in Figures 2A, 2C, and Figure 9C. **(A, B)** An RGC expressing either Bongwoori-R3 or GCaMP6f (Figure 10A or 10B) was patched in whole-cell configuration and stimulated electrically in current-clamp mode. At the bottom, there are electrophysiology signals from RGC, while the top shows where those signals were coming from. **(C, D)** The retina expressing Bongwoori-R3 was imaged while being electrically stimulated using either cathodal or anodal pulses. The bottom part displays how the retina was stimulated, while the top shows where the fluorescence response was distributed. “STIM” indicates the location of the stimulus. Stimulating cells either directly or indirectly can activate the cells themselves or nearby axon bundles, and this activation was seen at a specific location.

Furthermore, there is somatic, axonal, or indirect activation due to the effect of the stimulating electrode on the nearby axon track [26]. Somatic activation refers to the activation of the cell body near the electrode. Axonal activation refers to the activation of distant cells by stimulating the axon process. In patients with retinal implants, axonal activation stimulates peripheral ganglion cells, prolonging the perception and deteriorating the quality of the perception. Indirect activation refers to the concentric activation around stimulating electrodes. This leads to percept fading seen by retinal prosthetic implant patients. Among the three effects, a strategy that intensively stimulates cell bodies close to electrodes is effective in improving vision recovered through retinal prosthesis. Moreover, a pulse duration of less than 0.1 ms or more than 25 ms is required to prevent axonal activation [26]. Since Figure 9 of this paper used pulses with a duration of 1 to 10 ms, fluorescence signals in the retinal tissue appear to be network activity through axonal activation.

In conclusion, voltage and calcium sensors were applied onto photoreceptor-degenerated retinas to map retinal activity. The results shown here are the first to demonstrate that Bongwoori-R3 can report transmembrane potential changes in the retinal ganglion cells from the retinal explants. Many of the recently developed GEVIs restrict expression to the soma to reduce the background fluorescence in the recording. Such an approach does improve the signal to noise ratio by reducing the background fluorescence. However, Bongwoori-R3 activity can be detected without limiting its expression enabling signals in axons and axon bundles. When analyzing neuronal circuit activity, it will be of added value to not only detect where the signal is, but where it is not as well (Figure 7). Bongwoori-R3 has the capacity to facilitate dissecting the functional network between retinal neuron populations with the power of imaging connectivity via the activity of retinal ganglion cells.

## METHODS

### Animals

Animal experiments were performed in accordance with guidelines under protocols approved by the Institutional Animal Care and Use Committee of either (1) Seoul National University (SNU-210319-8), or (2) Korea Institute of Science and Technology (KIST-2021-01-018). Retinal degenerative mice (C57BL/6J strain, or C57BL/6J-*Pde6b*^*rd1-2J*^/J; #004766) were purchased from The Jackson Laboratory, USA and both male and female mice age from 6-week old were used.

### Intravitreal Adeno-associated Virus (AAV) Administration

Mice aged 6–8 weeks were anesthetized in an induction chamber (#941444, VetEquip, USA) with 3%–5% isoflurane in oxygen, and the subsequent virus injection was conducted under a LED stereoscopic microscope (SMZ 745, C-W10 ×B/22; Nikon, USA). The virus constructs encoding Bongwoori-R3 or GCaMP6f (constructs, AAV2/2.hSyn.Bongwoori-R3 and AAV2/1.Syn.GCaMP6f.WPRE.SV40; titers, 2.9×10^13^ and 1.2 ×10^13^ GC/mL; production, KIST Virus Facility, South Korea and Addgene #100837, USA) were separately loaded into a glass pipette (#4878; World Precision Instruments, USA).

Eyes were initially punctured with a beveled needle (NF36BV-2, NanoFil Needles, World Precision Instruments) in the *pars plana* site that is posterior from the *limbus* of the eye [41, 42], and injected with a volume of 2 *μ*L at a rate of 20 nL/s (Micro4, World Precision Instruments, USA) into the vitreous cavity (51730D, Stoelting, USA). At 2 weeks after injection, the retina was prepared for fluorescence imaging experiments.

### Tissue Preparation and Solutions

Mice were euthanized by cervical dislocation. Subsequent retinal preparation was conducted in Ringer’s solution—(in mM) 124 NaCl, 2.5 KCl, 2 CaCl_2_, 2 MgCl_2_, 1.25 NaH_2_PO_4_, 26 NaHCO_3_, 22 D-glucose—under a stereomicroscope (light source, SZ2-CLS; microscope objective lens, Olympus DF PLAPO 1 × -4; stereomicroscope system, SZX10; all by OLYMPUS, Japan). The eyeballs were dissected and immersed in the solution, and the retina was separated from the eyes (see section Retina Isolation) and mounted ganglion-layer up on membrane filter (HAWP04700, Merck, USA). The retina was then secured with slice anchors (SHD-27LH/10) in a bath chamber (RC-26G; both by Warner instruments, USA) under the one-photon fluorescent microscope. The solution was constantly bubbled with 95% O_2_ and 5% CO_2_ and perfused to the chamber at a rate of 1 mL/min, maintained at 34°C with an automatic temperature controller (TC-344B, Warner Instruments, USA).

### Retina Isolation

The cornea of the dissected eye was firmly held with forceps and incisions were made along the circumference of the cornea using scissors until it reached *pars plana*. The anterior part of the outer layer, iris, pigmented tissue, and lens were removed by using forceps. The optic nerve was removed by cutting out the base part attached to the eyecup using scissors. Then the outermost layer of the eyecup was cut open by making an incision starting from *pars plana*, passing the optic disc, and towards the *pars plana* on the opposite side.

Another cut was made at 90 degrees of the first incision line leaving four leaves so that the outermost layer of the eyecup could be carefully peeled off. The vitreous body was pulled away from the retina by using forceps while the retina was gently held with another pair of forceps without excessive pinching. Any leftover ciliary body, retinal pigment epithelium (RPE), or vitreous body was removed using forceps. The dissected retina was then flattened by making four halfway incisions from the retinal periphery towards the optic disc.

### Simultaneous Imaging and Electrophysiology

Upright one-photon fluorescence microscope system (Slicescope, Scientifica, UK) equipped with a 460 nm LED (UHP-Mic-LED-460, Prizmatix, Israel), GFP filter set (GFP-3035D-OMF, Semrock, USA), 10× (0.30 NA; Olympus, Japan) and 60× (1.00 NA, Olympus, Japan) water immersion objective lenses were used. LED light intensity for GFP excitation was illuminated as 1 mW/mm^2^ at a specimen plane. A highspeed color charge-coupled-device (CCD) camera was utilized by NeuroPlex camera software (RedShirtImaging, USA). 80×80 pixels’ fluorescent images were captured at a rate of 1 kHz. The total acquisition time of imaging and recording was matched.

Membrane potentials were acquired through either tight-seal whole-cell or loose-seal cell-attached patch-clamp recording. The loose patch was made without excising the cell membrane, unlike a tight patch. Patch pipettes were fabricated from borosilicate glass capillaries (1B150F-4, World Precision Instruments, USA) on a micropipette puller (P-97 Sutter Instrument, USA). The pipettes had a resistance of 5– 15 MΩ when filled with the normal internal solution—(in mM) 120 K-aspartate, 4 NaCl, 4 MgCl_2_, 1 CaCl_2_, 10 EGTA, 3 Na_2_ATP, 5 HEPES—and were held by a pipette holder containing an Ag/AgCl electrode (Molecular Devices, USA) and mounted on a motorized micromanipulator (Scientifica, UK). Whole-cell and cell-attached patches employed an amplifier (Multiclamp 700B) coupled to a digitizer (Axon Digidata 1550B), which was configured on the software (pClamp 10; all by Molecular Devices, USA). Signals were digitally sampled at 10 kHz in the bridge mode between current injection and recordings. Before electrophysiology, inner limiting membranes were carefully torn by glass pipettes to record underneath ganglion cells of interest.

### Electrical Stimulation

For intracellular stimulation, current pulses (whole-cell patch, 100 pA, 100 or 200 ms; cell-attached patch, 50 pA, 200 ms) were delivered into ganglion cells through patch pipette in current-clamp mode, and voltage pulses (wholecell patch; -120 to 120 mV, 200 ms, or -150 to 150 mV, 300 ms; holding potential, -70 mV) were delivered in voltage-clamp mode. In each cell, patch experiments were repeated three times or two times with current-clamp or voltage-clamp, respectively. For field stimulation, either anodal or cathodal current pulses (0.2 to 8 mA, 0.1 to 10 ms) were generated by a constant stimulator (DS3, Digitimer Ltd, England) and delivered onto the retina near ganglion cells of interest through bipolar microelectrode (#30202, FHC, USA). Both intracellular and extracellular stimulations were triggered to start by pClamp software and Digidata 1550B digitizer.

### Data Analysis

Electrophysiological data were inspected using Clampfit 10.7 software (Molecular Devices, USA) and imaging data using NeuroPlex version 10.2.0 (RedShirtImaging, USA) and ImageJ (National Institutes of Health, USA). Data were analyzed by Excel 2016 (Microsoft, USA) for summarized and transformed data and by Matlab (Mathworks, USA) for computation, display purposes and statistical tests. Statistical differences were tested with relevant functions in Matlab. Graphical illustration was created with BioRender.com.

Electrical signals were analyzed through Clampfit software. The value for an amplitude of membrane potentials was quantified relative to the first value. Electrical signals were synchronized to optical traces through NeuroPlex. Optical signals were processed through NeuroPlex software. Optical traces data were applied Gaussian filter at a low pass of 50 Hz unless otherwise noted and displayed fluorescence amplitude over time. The value for fractional fluorescence change (ΔF/F) was calculated as dividing the difference between a peak height and a baseline by the fluorescence intensity at rest in the region of interest (ROI), which is in the form of ΔF/F = [(F_X_ – F_0_) / F_0_] × 100. In Neuroplex, the function for frame subtraction determines the difference in fluorescence values of all pixels acquired in two frames of the holding potential and the stimulation periods. Further analysis for both optical and electrical signals was determined and calculated through Matlab. The parameter for a lag time was defined as the delay of the optically recorded first AP to the electrically recorded first AP. The parameter for fall time was defined as a time difference between the optical peak and baseline signal. The parameter for decaying fractional value (ΔF/F) was defined as a difference of fluorescence intensity from the first 100 frames to the last 100 frames.

Movie clips were made based on imaging stacks and field of view (FOV) images. As one component, the FOV fluorescence image was captured as a still image. As the other component, imaging stacks in which pixels represent fluorescence changes (ΔF) were obtained within NeuroPlex. For spatial processing, the imaging stack was low-pass filtered using a 3 × 3 mean kernel and iterated 3 times, and the movie was scaled by min-max of all diodes. The imaging stacks were binned every 10 images to make movies within ImageJ. Additional conditioning was processed through Adobe Premiere Pro (Adobe System, USA). The playback speed was adjusted depending on the length of the movies to 2.75×, 1×, 1.75× and 1.75× in Movies 1–4, respectively.

### Statistical Analysis

Prior to statistical comparison, Jarque-Bera method was used to test null hypothesis that the data is normally distributed with unspecified mean and standard deviation [42]. When this normality test cannot be rejected, two-sample *F*-method was used to test null hypothesis that two data have same variances [42]. When this variance test cannot be rejected, *p* values were computed with two-sample Student’s *t*-test. Statistical significance was determined by α = 0.05. Numerical value ranges were given as the mean ± SD.

## ASSOCIATED CONTENT

### SUPPORTING INFORMATION

The Supporting Information is available free of charge at …

Movies of imaging action potentials from either the GEVI, Bongwoori-R3 or the GECI, GCaMP6f expressing retinal ganglion cells of *rd1* blind mice in response to current-clamp steps (MP4); Movies of imaging action potentials from the GEVI, Bongwoori-R3 expressing retina tissue in *rd1* blind mice during electrical stimulation with epi-retinally positioned microelectrode (MP4)

## AUTHOR INFORMATION

### Author Contributions

Baker BJ and Song YK supervised this study; Jung Y acquired and analyzed data; Jung Y, Lee S and Rhee JK drafted the manuscript; Rhee JK produced AAV vectors; Lee CE assisted to prepare movies; All authors contributed to revising the manuscript or figures.

## ACKNOWLEDGMENT

The author would like to thank Prof. Jiayi Zhang’s laboratory at Fudan University in China for technical help. The author also would like to thank Dayoung Lee from Seoul National University in South Korea for helpful comments on the earlier version of the manuscript.

## ABBREVIATIONS

AP: action potential
FOV: field of view
ΔF/F: fractional fluorescence change
GECI: genetically-encoded calcium indicator
GEVI: genetically-encoded voltage indicator
RGC: retinal ganglion cell
ROI: region of interest.

## For Table of Contents Only

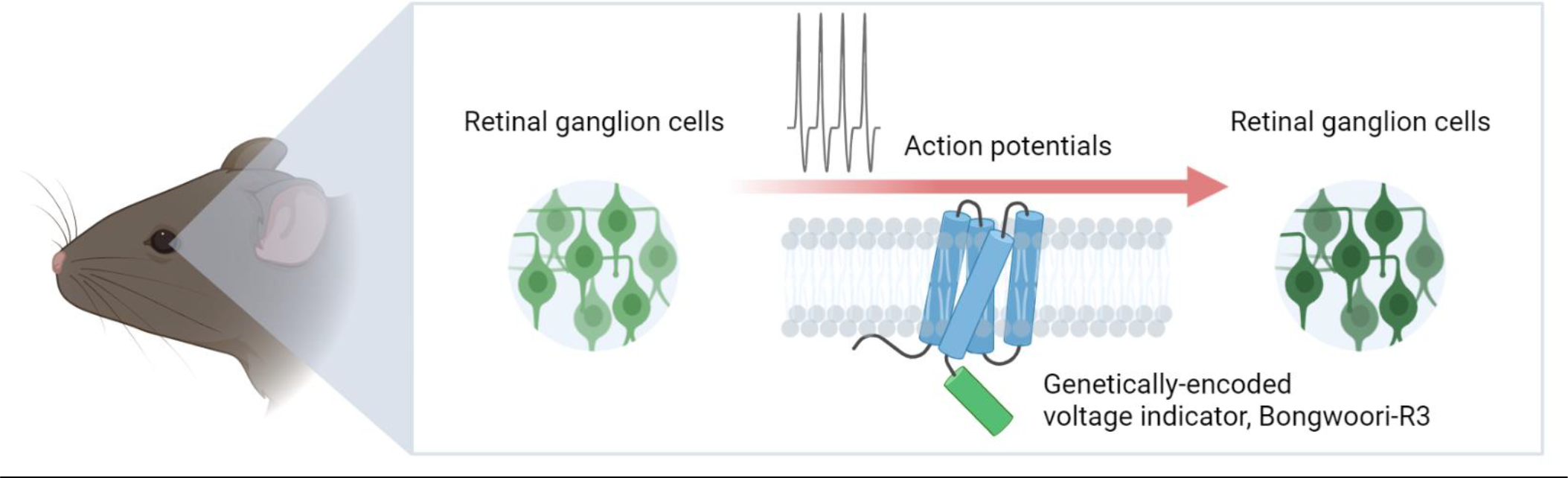

