## Supporting Information for "Imaging Electrical Activity of Retinal Ganglion Cells with Fluorescent Voltage and Calcium Indicator Proteins in Retinal Degenerative *rd1* Blind Mice"

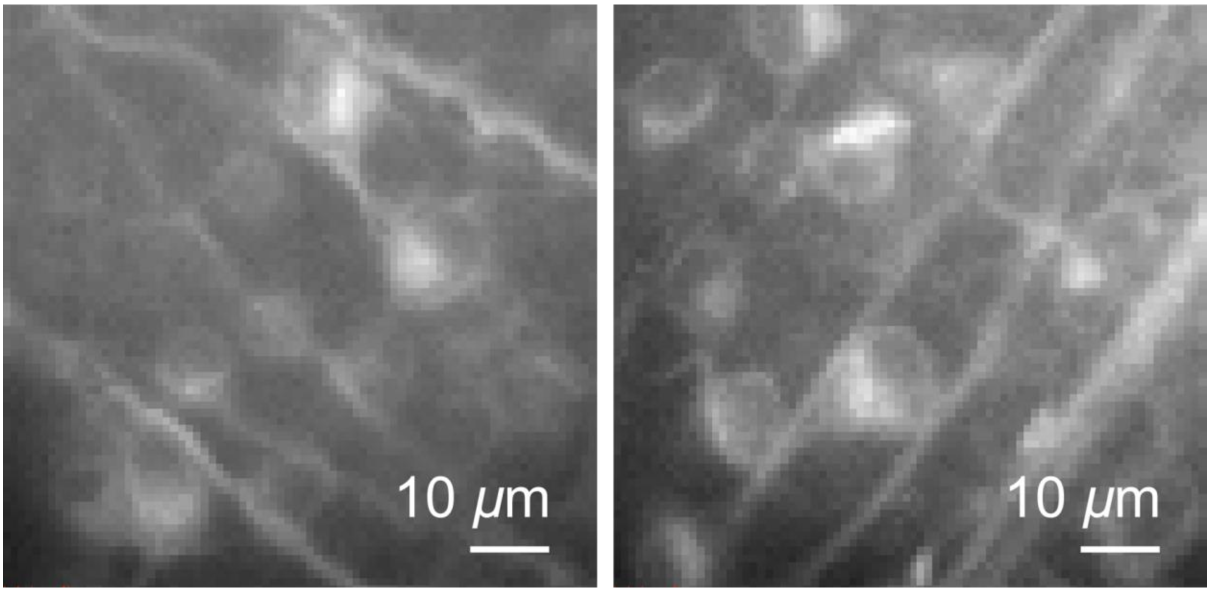


**Figure S1. Fluorescence images of retinas from *rd1* blind mice.** Retinal ganglion cells (RGCs) and their axon bundles that express the voltage indicator, Bongwoori-R3.

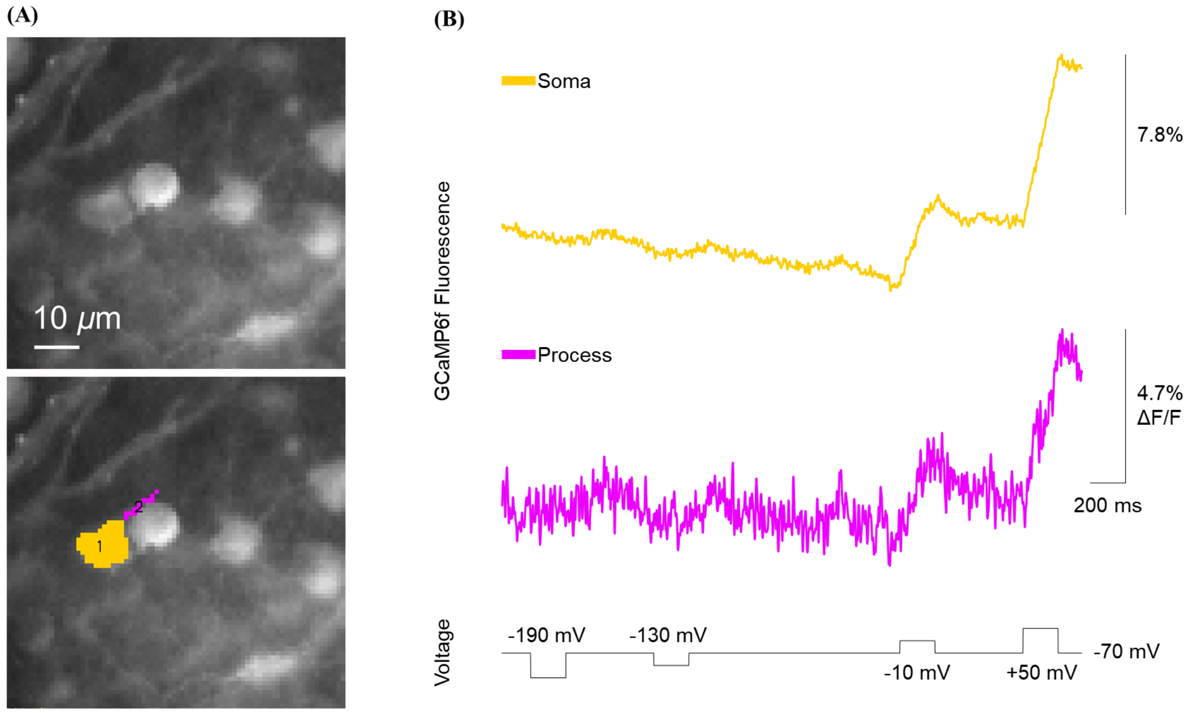


**Figure S2. Electrophysiological and fluorescence signals from the GECI, GCaMP6f expressing retinal ganglion cells of *rd1* blind mice in response to voltage-clamp steps.** Fluorescence and electrophysiology signals were recorded at the same time. The fluorescence data was averaged over 16 trials, with each trial being sampled at a rate of 1 kHz. The average was then filtered using a Gaussian filter that removed any frequencies above 20 Hz. **(A)** The top image shows the resting fluorescence of retinal ganglion cells that express the calcium indicator, GCaMP6f. The bottom image shows the pixels of interest corresponding to the retinal ganglion cell and its process. The fluorescence was measured by averaging the pixels of interest in the image. This is then shown as color traces, which show how the fluorescence levels change over time. **(B)** The voltage of Soma (yellow trace) was regulated to match the voltage changes in the step protocol at the bottom (black trace). The step protocol had an amplitude of -120, -60, 60, and 120 mV, a duration of 200 ms, and a holding potential of -70 mV. The calcium sensor, GCaMP6f exhibited slow fluorescence changes and was detected in the processes of retinal ganglion cells.

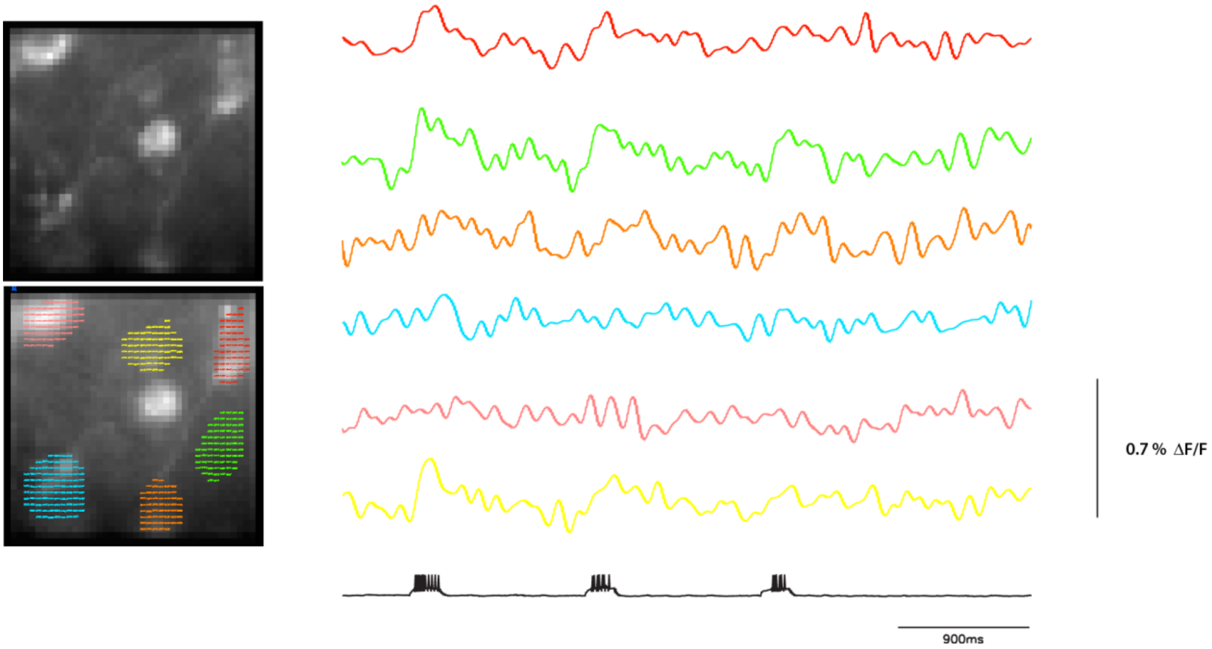


**Figure S3. Communication among retinal ganglion cells in *rd1* blind mice.** Retinal ganglion cells (RGCs) were imaged using a fluorescent protein called Bongwoori-R3 at a rate of 3 kHz. At the same time, the cell was also recorded using current-clamp mode at a rate of 10 kHz. These signals were displayed in either color or black. The electrophysiology and fluorescence signals were both recorded from each single trial. The fluorescence traces were then Gaussian filtered at a low pass setting of 10 Hz or 30 Hz. This figure was produced by the same data as in Figure 5. **(Left)** The top image shows the resting fluorescence of retinal ganglion cells that express the Bongwoori-R3 protein. The bottom image shows specific pixels representing these cells of interest. The fluorescence emitted from the cells was obtained by averaging the pixels of interest in the image. This is then shown as color traces, which show how the fluorescence levels change over time. **(Right)** When electrically stimulated (Figure 5; green ROI), the retinal ganglion cells exhibited fluorescence signals in response. The stimulation consisted of applying an electrical current of 100 pA for 200 ms using the whole-cell patch configuration. Red, green, yellow ROIs exhibited a fluorescence response to the first pulse, whereas green ROI responded to all three pulses. See also Figure 5.

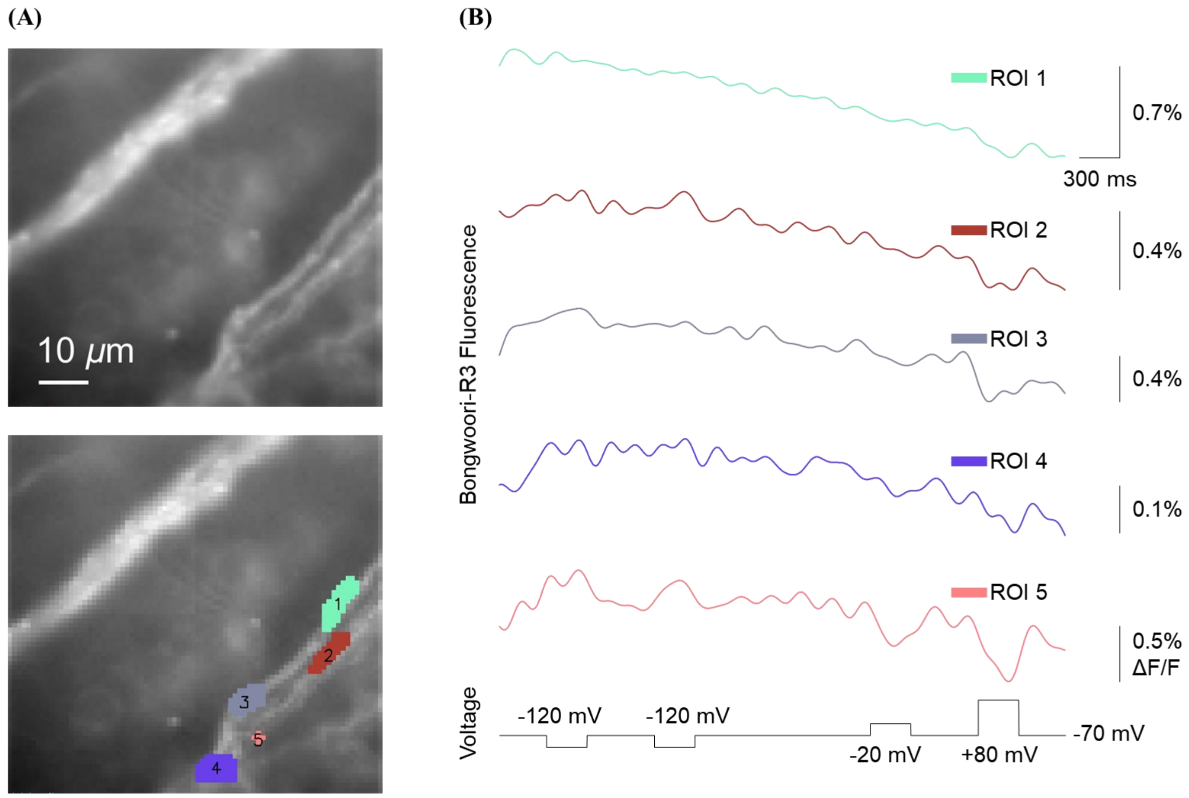


**Figure S4. Tracing the voltage signal along an axon bundle.** Retinal ganglion cells (RGCs) were imaged using a protein sensor called Bongwoori-R3 and simultaneously recorded using voltage-clamp. The fluorescence data was averaged from 16 trials, each sampled at 1 kHz. The data was then filtered using a Gaussian filter with a low pass set at 5 Hz. The fluorescence signal is presented in color. This figure was produced by the same data as in Figure 6. **(A)** The upper image shows the resting fluorescence of retina expressing Bongwoori-R3. The lower image shows specific pixels of interest that correspond to different parts of axon bundles. **(B)** The fluorescence was obtained by averaging the pixels of interest. This is then shown as color traces. The voltage of Soma 1 (Figure 6B; red trace) was controlled to follow the step protocol at the bottom (black trace), which had an amplitude of -50, -50, 50, and 150 mV, a duration of 300 ms, and a holding potential of -70 mV. Applying voltage steps to Soma 1 (Figure 6B; red trace) caused a fluorescence signal to appear in ROI 5 (pink trace), but not in the lower axon bundle (ROI 4). Distinct voltage signals were detected along different regions of the axon bundle. ROI 1–3 exhibited small but obvious large depolarization signals. ROI 2 also showed a slight hyperpolarization signal, whereas ROI 5 showed a clear hyperpolarization signal. See also Figure 6.

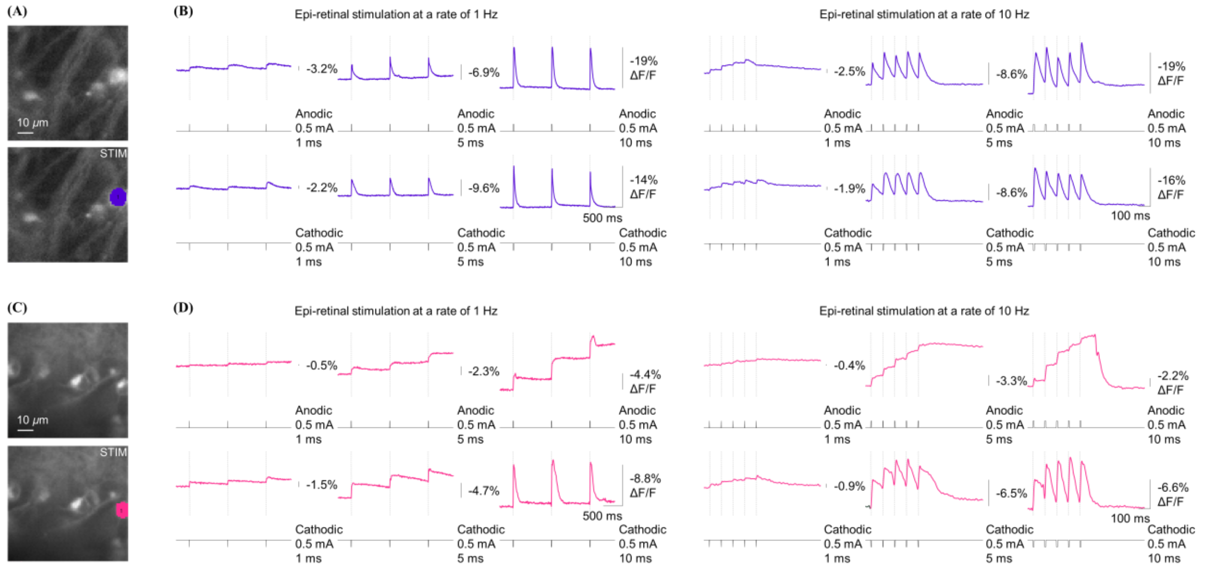


**Figure S4. Fluorescence signals from the GEVI, Bongwoori-R3 of retina tissue in *rd1* blind mice during electrical stimulation with epi-retinally positioned microelectrode.** While the retina was being stimulated with a microelectrode, fluorescence signals were acquired from a single trial at a sampling rate of 1 kHz. The signals are shown in color. **(A, C)** The top image shows the retina's resting fluorescence that expresses the Bongwoori-R3. The bottom image shows the specific pixels that were used for analysis. A stimulating microelectrode was located in the upper right corner, outside the field of view. This is indicated by the "STIM" label. The fluorescence from the RGC was measured by averaging the pixels of interest in the image. This is then displayed as color traces, which show how the levels of fluorescence changes over time. **(B, D)** When the retina was electrically stimulated (black trace), it resulted in the generation of fluorescence signals (color trace). The stimulus was indicated on the right side of each trace (amplitude, 0.5 mA; duration, 1, 5, 10 ms; rate, 1, 10 Hz). For reference, the vertical dash lines indicate the start of the stimulus pulse. How retinal cells respond to different stimulus frequencies and parameters was compared by measuring their fluorescence response. This fluorescence represents alterations in the membrane potential of the cells.

**Movie S1:** A retinal ganglion cell (RGC) expressing Bongwoori-R3 was patched in whole-cell configuration and stimulated electrically in current-clamp mode. (amplitude, 100 pA; duration, 200 ms).

**Movie S2:** A retinal ganglion cell (RGC) expressing GCaMP6f was patched in whole-cell configuration and stimulated electrically in current-clamp mode. (amplitude, 2000 pA; duration, 200 ms).

**Movie S3:** A retina expressing Bongwoori-R3 was imaged while being electrically stimulated using cathodal pulses (amplitude, 2 mA; duration, 10 ms; rate, 10 Hz).

**Movie S4:** A retina expressing Bongwoori-R3 was imaged while being electrically stimulated using anodal pulses (amplitude, 2 mA; duration, 10 ms; rate, 10 Hz).
